# Recent visual experience reshapes V4 neuronal activity and improves perceptual performance

**DOI:** 10.1101/2023.08.27.555026

**Authors:** Patricia L. Stan, Matthew A. Smith

## Abstract

Recent visual experience heavily influences our visual perception, but how this is mediated by the reshaping of neuronal activity to alter and improve perceptual discrimination remains unknown. We recorded from populations of neurons in visual cortical area V4 while monkeys performed a natural image change detection task under different experience conditions. We found that maximizing the recent experience with a particular image led to an improvement in the ability to detect a change in that image. This improvement was associated with decreased neural responses to the image, consistent with neuronal changes previously seen in studies of adaptation and expectation. We found that the magnitude of behavioral improvement was correlated with the magnitude of response suppression. Furthermore, this suppression of activity led to an increase in signal separation, providing evidence that a reduction in activity can improve stimulus encoding. Within populations of neurons, greater recent experience was associated with decreased trial-to-trial shared variability, indicating that a reduction in variability is a key means by which experience influences perception. Taken together, the results of our study contribute to an understanding of how recent visual experience can shape our perception and behavior through modulating activity patterns in mid-level visual cortex.

## Introduction

Prior knowledge biases our perception of incoming sensory input. This can occur through a mixture of top-down cognitive factors and an accumulation of sensory experience that governs how our sensory systems process and interpret incoming information. From moment to moment, recent experience with a visual input has both neural and cognitive impacts. On the order of milliseconds to minutes, repetition of a stimulus leads to modulation or adaptation of sensory neuron responses (Kohn, 2007; Solomon and Kohn, 2014; Webster, 2015) and is linked to perceptual changes such as visual aftereffects (Clifford et al., 2007; Webster, 2015; Weber and Fairhall, 2019). Repeated exposure can also create an expectation of an upcoming sensory event. An increase in expectation, or the probability of a sensory event occurring, both shapes our neural activity and improves perceptual abilities (Summerfield and de Lange, 2014; de Lange et al., 2018) such as more rapidly or accurately detecting a change in incoming visual input (Wyart et al., 2012; Pinto et al., 2015; Stein and Peelen, 2015). Given that recent sensory experience can have a profound impact on perception and behavior, understanding the neural mechanisms by which our brains incorporate sensory contexts and experiences is critical.

The impact of recent visual experience has been studied on various timescales. On the shortest timescale (i.e. milliseconds to seconds), the most common are studies of adaptation. The prevailing finding under the umbrella of adaptation effects is that of repetition suppression - a reduction in response following the repeated presentation of a stimulus (Kohn, 2007; Solomon and Kohn, 2014; Webster, 2015; Weber et al., 2019). Recent experience on longer timescales (i.e. minutes or across trials) can also impact neurons and behavior. Studies of expectation use repeated presentation of a stimulus (Summerfield et al., 2008; Kaliukhovich and Vogels, 2010; Meyer and Olson, 2011; Ramachandran et al., 2016, 2017; Kumar et al., 2017; Pajani et al., 2017) or cues (Summerfield and Koechlin, 2008; Egner et al., 2010; Kok et al., 2012; John-Saaltink et al., 2015; Amado et al., 2016; Bell et al., 2016; Davis and Hasson, 2018; Dunovan and Wheeler, 2018; Rungratsameetaweemana et al., 2018) to modulate the probability of an event occuring and are associated with a suppression of activity with greater expectation (Summerfield and de Lange, 2014; de Lange et al., 2018; but see Rao et al., 2012; Feuerriegel et al., 2021). However, the relationship between within-trial (short time scale) and across-trial effects of recent experience remains unclear (Summerfield et al., 2008; Kaliukhovich and Vogels, 2010, 2014; Kovacs et al., 2013; Grotheer and Kovács, 2014, 2015; Kronbichler et al., 2018; Vinken et al., 2018).

Although there is a large literature separately reporting perceptual and neural effects of recent experience, only a small number of adaptation and expectation studies directly link neural measurements to the behavioral consequences of recent visual experience (Dragoi et al., 2002; McDermott et al., 2010; Kok et al., 2012; Wissig et al., 2013; Bell et al., 2016; Jin et al., 2019). Instead, most task paradigms investigating recent experience along with neural measurements involve merely the passive viewing of images or tasks that do not directly link recent visual experience with improvements in psychophysical performance (Summerfield et al., 2008; Egner et al., 2010; Kaliukhovich and Vogels, 2010; Meyer and Olson, 2011; Amado et al., 2016; Ramachandran et al., 2016, 2017; Kumar et al., 2017; Kaposvari et al., 2018; Richter et al., 2018; Ghodrati et al., 2019; Vergnieux and Vogels, 2020; Nigam et al., 2023). This relative lack of joint neural and behavioral measurements of recent experience leaves a gap in our understanding of these phenomena. In addition to changes in individual neuron responses, it is possible that the effects of experience are mediated by changes in correlated activity across cortical populations of neurons. Theoretical work (Shadlen and Newsome, 1998; Abbott and Dayan, 1999; Averbeck et al., 2006; Moreno-Bote et al., 2014; Sharpee and Berkowitz, 2019; Bartolo et al., 2020) and experimental studies of cognitive factors such as attention (Cohen and Maunsell, 2009; Mitchell et al., 2009; Herrero et al., 2013; Ruff and Cohen, 2014a, 2019; Ni et al., 2018; Snyder et al., 2018) indicate that changes in the trial-to-trial correlated activity of a population of neurons can improve stimulus discriminability. However, few studies have assessed the impact of recent visual experience on these population level interactions, leaving many unanswered questions about how recent experience shapes interactions between visual neurons.

Our goal was to investigate the neural impact of recent experience on visual cortical area V4 during a task where greater recent visual experience could improve visual perception. We designed a natural scene change detection task that used stimulus probability to maximize and minimize the amount of recent experience with a particular image, and assessed the effects of experience on V4 neuron firing rates and population activity. We found that maximizing recent experience led to a suppression of V4 neural activity that was correlated with an improvement in behavioral performance. This decrease in activity resulted in an increase in signal separation between the image and the target (the changed image) to be detected. Additionally, we found that greater experience was associated with reduced trial-to-trial shared variability among neurons, and this reduction was related to correct trial outcomes. Our work thus establishes neural correlates of recent visual experience, at the level of individual neurons and populations, that appear to underlie the behavioral improvements associated with recent experience.

## Methods

### Experimental Subjects

All experimental procedures were conducted in accordance with the United States National Research Council’s Guide for the Care and Use of Laboratory Animals and approved by the Institutional Animal Care and Use Committee of Carnegie Mellon University. Data from two male rhesus macaque monkeys were included in this study. Prior to the start of behavioral training, monkeys were surgically implanted with a titanium headpost using aseptic techniques under isoflurane anesthesia. Once monkeys showed proficiency in the task, we surgically implanted a 100-electrode Utah array (Blackrock Microsystems) in left hemisphere V4.

### Electrophysiological recordings

Signals from the microelectrode array were bandpass filtered (0.3Hz to 7.5 kHz), digitized at 30 kHz, and recorded by a Grapevine system (Ripple Neuro). Waveform segments exceeding a threshold (-3 times the root mean square noise on each array channel) were saved for offline processing. Following data collection, we performed spike sorting in two stages. First, waveforms were automatically sorted as either spikes or noise using a neural network classifier developed in our laboratory (Issar et al., 2020). Second, we manually spike-sorted all of the waveforms that passed the first stage using custom software developed in our laboratory (Kelly et al., 2007) using the time-voltage waveforms, interspike interval distribution, and waveform shape characterized by principal component analysis. The output of this spike sorting process included both well-isolated single units and small multi-unit groups with similar waveforms. We refer to each of these as a “neuron”.

### Receptive field mapping

All visual stimuli were shown on a luminance-corrected CRT monitor positioned 57 cm away from the monkey’s eyes. Monocular eye position and pupil diameter were constantly monitored using an infrared camera (EyeLink 1000). We mapped receptive fields (RFs) of the V4 neurons to determine appropriate size and location for the images used in the visual change detection task. Sinusoidal gratings of 4 possible orientations were displayed one at a time in a grid of positions that covered the likely RF area based on the anatomical location of the implant. For monkey PE, 1.9 degree gratings covered an area 1.9 degrees above fixation to 11.0 degrees below fixation and 1.9 degrees to the left of fixation to 14.7 degrees to the right of fixation. For monkey RA, 0.9 degree gratings covered an area 2.8 degrees above fixation to 7.4 degrees below fixation and 1.9 degrees to the left of fixation to 7.4 degrees to the right of fixation. We identified the center and extent of each neuron’s receptive field by analyzing the mean response to gratings presented at each location. The size and position of images for the change detection task were chosen to cover the aggregate of the V4 receptive fields recorded by the array. For Monkey PE, the image was 10.6 degrees in diameter and centered at 4.4 degrees below and 6.2 degrees to the right of fixation. For Monkey RA, the image was 8.0 degrees in diameter and centered at 3.8 degrees below and 3.8 degrees to the right of fixation.

### Visual stimulus selection

We chose to use natural images (as opposed to oriented gratings or other parameterized images) in order to capture more diverse and naturalistic visual responses from our populations of neurons. To this end, the precise symbolic content of the image (i.e. how likely our subjects are to encounter the scene in their natural environment) was less important than ensuring that the images selected drove strong and diverse responses in our neuronal population. Natural images were acquired from Yahoo Flickr Creative Commons 100 Million Dataset and consisted of 112x112 pixel images of a variety of natural scenes and objects. To select the images used for daily experiments, we used a stimulus selection paradigm (Cowley et al., 2017) to identify images that drove strong and diverse responses in the particular V4 population being recorded. We ran multiple recording sessions (6 for monkey PE, 2 for monkey RA) where in each session we presented 1200-2000 new natural images. Monkeys performed an active fixation task while passively viewing sequences of 6 (Monkey PE) or 8 (Monkey RA) images. On each trial, 6 or 8 images were pseudo randomly selected from the larger pool for that day. Each image was displayed for 100 ms with a 100 ms inter-stimulus interval. Each image was repeated 10-24 times, depending on the number of images and duration of that session.

After gathering image responses, the 600 images eliciting the strongest population firing rate (average across neurons) were selected. We then manually curated the group of images to eliminate those that were either lacking in color (e.g. a grayscale image) or orientation information (e.g. a blue sky) which would have made it nearly impossible to detect a color or orientation change, respectively. For the remaining 300-400 images, we used principal component analysis (PCA) on the average response to each image to identify the principal components (PCs) that explained the greatest response variance. We projected the neural responses into this PCA space. To select groups of 10 images for each session of the main change detection task, we used an objective function that favored large responses that were far away from each other in PCA space.

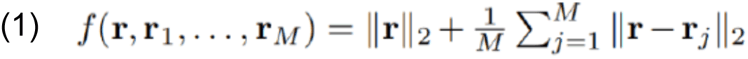

Here **r** is a vector of responses for each neuron. *M* is the number of images. The term 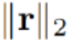

maximizes the responses of the group of neurons, while the term 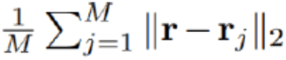 maximizes the Euclidean distance between **r** and **r**1,…,**r**_M_. Using this method, we were able to select new images for each session that would result in strong and diverse responses in our recorded V4 neurons.

### Natural image change detection task paradigm

Subjects performed a natural image change detection task where the subject’s goal was to determine and report when a flashing natural image had changed (Fig 1). Following a fixation period (300 ms), a natural image (the “sample”) was displayed on the screen for 300 ms, followed by a gray screen inter-stimulus interval (lasting 250-350 ms, chosen randomly from a uniform distribution within that range). The image was repeatedly presented (“flashed”) a variable number of times, with a fixed probability (34% for Monkey PE and 36% for Monkey RA) that the target would appear on each flash after the original sample. This created a flat hazard function to discourage guessing. The sample image was fixed for each trial and remained the same across flashes until the target (a changed version of the sample image) appeared. Once the target was presented, the subject had 400 ms to saccade to the target and receive a liquid reward. Subjects had to detect a change in either orientation or color of the sample image on that trial. The change type was fixed for the entirety of a session, and we alternated orientation and color sessions. There were 4 difficulties of targets which were determined by a rotation in orientation or L*a*b color space. Difficulties were titrated to obtain comparable average performance for both change types. To additionally encourage constant behavior within a trial, the reward was adjusted based on the number of flashes, such that longer trials resulted in a higher reward. The ramping of reward was dependent on the animal and set at a value that encouraged a flat false alarm rate over the duration of each trial. This was done to promote relatively constant attention and motivation throughout the trial.

**Figure 1.**
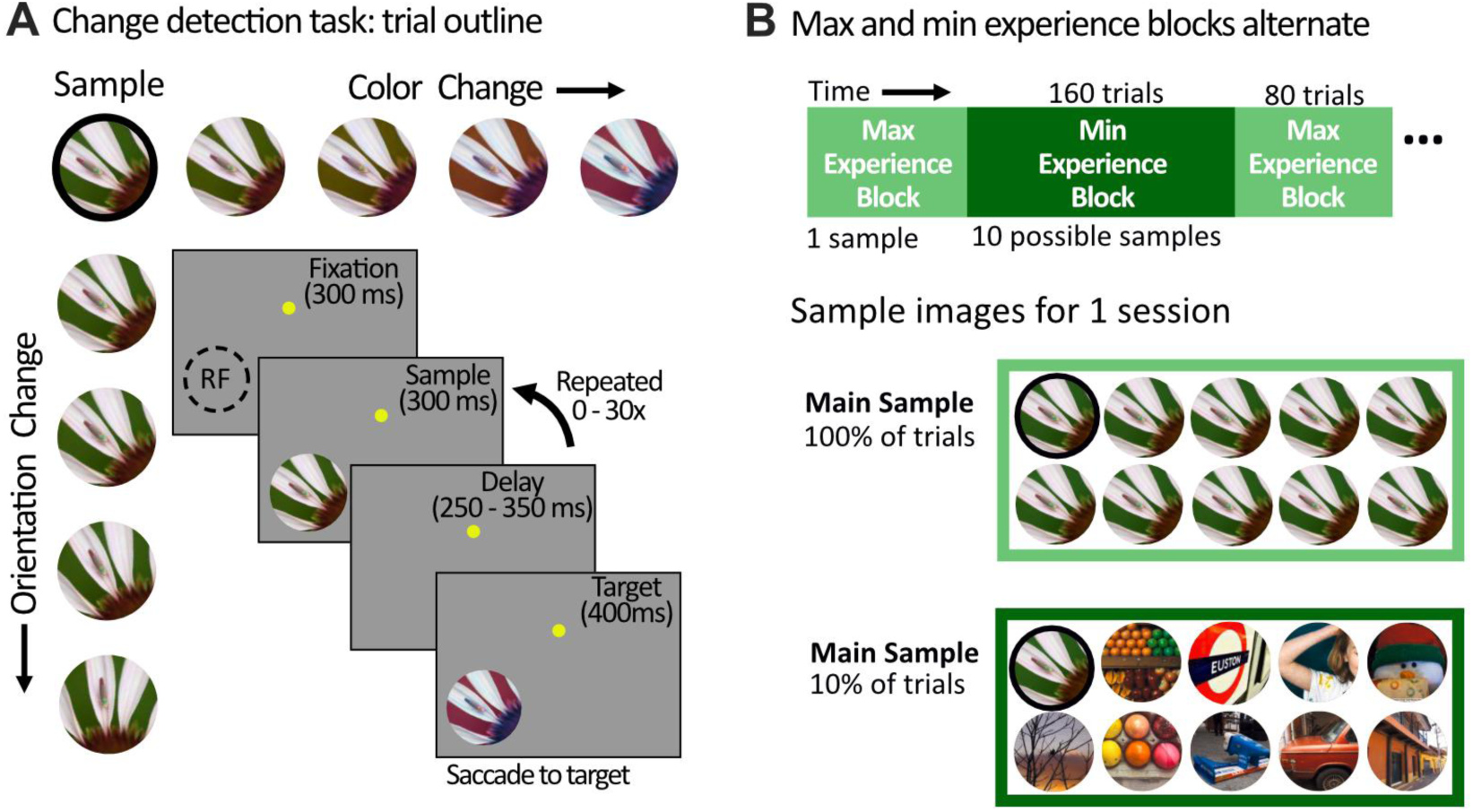
Modulating stimulus probability to create different levels of recent visual experience in a natural image change detection task. A) Natural image change detection task outline. A trial began when the monkey fixated on the yellow dot. On each trial, a sample image was chosen and repeatedly flashed on the screen in the area of the V4 receptive fields. On a flash unknown to the monkey (fixed probability), a feature of the image changed, and the monkey was rewarded for a saccade to the changed image (the target). Targets could be one of 4 difficulties. Sessions of color and orientation changes were alternated. B) Stimulus probability was used to create max and min experience conditions. Top; Max and min experience blocks of trials alternated throughout the session. Bottom; Example images are shown for each condition for one session. To create max and min experience conditions, we modulated the stimulus probability of a particular image (the “main sample”). In max experience blocks, the main sample was the sample image on 100% of trials. In min experience blocks, the main sample was the sample image on only 10% of trials, and the other 90% of trials used one of 9 different images as the sample image (each shown on 10% of trials). The main sample image for an example session is circled in black.

**Figure 2.**
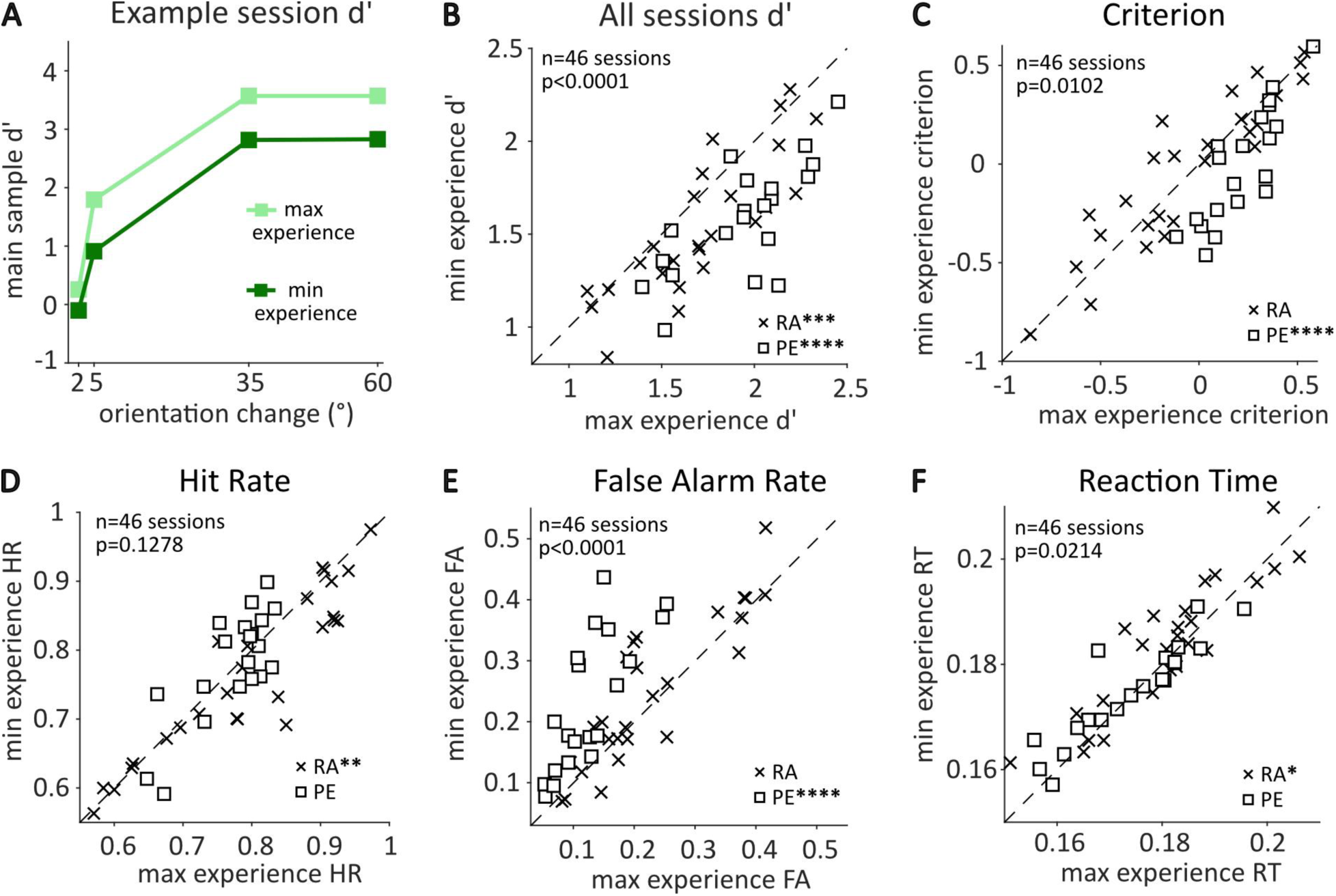
Greater recent experience with an image led to improved perceptual performance on a natural image change detection task. A) The behavioral performance for the main sample image in one session showed an improvement in discriminability (*d’)* for detecting each change difficulty in the max experience condition compared to the min experience condition. B) Comparison of *d’* across sessions for two monkeys indicated a robust improvement in discriminability in the max experience condition (paired t-test, p<0.001 for RA, p<0.0001 for PE). For each session, *d’* (collapsed across target difficulties) for the main sample was determined in each condition. C) Criterion was significantly higher in the max experience condition for monkey PE (paired t-test p<0.0001) but not monkey RA (p=0.4086). D) The difference in hit rates between max and min experience across sessions was significant for Monkey RA (top, paired t-test p<0.01) but not monkey PE (p=0.4526). E) The difference in false alarm rates between max and min experience conditions across sessions was significant for monkey PE (paired t-test p<0.0001) but not monkey RA (p=0.0698). F) Reaction time was calculated across all correct trials for each session. There was a significant decrease in reaction time in the max experience condition for monkey RA (paired t-test, p<0.05) but not monkey PE (p=0.3393).

To create differences in recent visual experience, we manipulated the probability of a particular sample image (the “main sample”) appearing on a given trial. In blocks of trials with maximal recent experience (“max experience”), the main sample was used on 100% of trials, meaning the subject always had to detect a change in the main sample. In blocks of trials where we minimized recent experience (“min experience”), the main sample was used on only 10% of trials, and 9 other images were used for the remaining 90% (each image appearing on 10% of trials). 10 new images were chosen each day (see Visual stimulus selection), one of which was selected as the main sample for that day. In the min experience blocks, images were pseudo randomly chosen for each trial. Max experience blocks contained 80 completed trials, and min experience blocks contained 160 completed trials (16 trials of each of the 10 images). Importantly, block transitions were triggered by 80 or 160 completed trials (i.e. either a correct detection of the change, or a failure to notice the image had changed). Broken fixations, false alarms, or failure to engage with the task did not count towards this trial count. Therefore, the time it took to complete a block was quite variable, meaning the animals could not use specific timing cues to determine block switches. Block transitions were not cued, so subjects had to rely on visual experience for knowledge of the block identity. Max and min experience blocks alternated throughout the day (a total of 6-16 blocks were completed each session), and which condition was the starting block was alternated for each subsequent session.

To understand how manipulating stimulus probability in this way could produce a difference in behavioral performance, it is helpful to consider the following example. In a max experience block, following 5 trials where the main sample was used on every trial, at the start of fixation on the 6th trial the observer would likely expect to see the main sample again given their recent visual experience. Given that the change detection needs to be made quickly and often with few flashes of the sample image, expecting to see a particular image could confer an advantage in detecting the change. In a min experience block, following 5 trials where a different image was the sample on each trial, at the start of fixation on the 6th trial the observer would not know which image to expect (it could be any one of the 10). In this case, if the main sample appeared on the 6th trial, the expectation would be much lower than in the max experience scenario. Our analyses thus focused on an identical visual input (the main sample) when it occurred in two different task contexts (maximizing and minimizing recent experience). Importantly, the min experience block contained 10 images of equal probability, therefore the main sample was not unexpected or surprising, allowing us to assess the effects of recent experience without the confound of surprise.

### Behavioral analysis

There are 3 different trial outcomes that we used for measuring behavioral performance: corrects, misses, and false alarms. A “correct” trial was one in which the monkey succeeded in detecting a change in the sample image, made a saccade to the target within the appropriate time window, and received a reward. A “miss” trial was one in which the sample changed, but the monkey failed to detect the change and continued fixating until the target display ended. A “false alarm” trial was any time the monkey made a saccade toward a flashed sample image to report a change, but the sample image had not yet changed. Because the change could never occur on the first flash, a false alarm could only occur on flash 2 or later. The fourth true outcome in a two-alternative forced choice task, a “correct withhold”, also occurred in our task. In each trial if the animal maintained fixation through flashes of the sample image, those flashes were considered “correct withholds”. Because every trial ended with a change if the animal maintained fixation long enough, the trial itself could not terminate in a correct withhold.

We quantified several measures of behavioral performance for the main sample separately for max and min experience conditions. Blocks of the same condition were combined. When determining the performance for the main sample in the min experience condition, we only included trials of the main sample (and excluded trials of the other 9 stimuli). Hit rate was calculated as HR = (# correct trials)/(# of correct + miss trials). False alarm rate was calculated as FAR = (# of false alarm trials/total number of opportunities to false alarm). The opportunities to false alarm were the sum of total flashes where the subject could have looked at the image (leading to a false alarm trial) but did not (i.e., correct withholds). Using hit rate and false alarm rate, we calculated two measures from signal detection theory: d-prime (*d’*) and criterion. *d’* is a measure of sensitivity that allows us to measure a subject’s ability to detect a signal. *d’* = z(HR)-z(FAR) where z(HR) and z(FAR) are the z transforms of hit rate and false alarm rate. Criterion (c) is a measure of the bias in reporting a signal and is calculated as c= -0.5(z(HR) + z(FAR)). Additionally, we determined reaction times for correct trials, calculated as the time between the target onset and the moment the subject’s eyes reached the target.

Relative difficulty was calculated as the average performance (*d’*) for the other 9 images in the min experience condition minus the *d’* of the main sample in the min experience condition. Positive values indicated that the performance for the main sample was worse than the other images, therefore the main sample was on average a more difficult image in which to detect a change. Negative values indicated that the performance for the main sample was better than the other images, therefore the main sample was on average a less difficult image in which to detect a change.

### Stimulus probability control task

We collected 10 sessions of data from 1 monkey (RA) on a modified task design to determine if the number of images in each block had an effect on behavioral performance. The overall trial structure of the task was the same as described above, the only difference was in the number and frequency of images shown in the different conditions (Fig 3). The min experience condition described above remained unchanged, we simply renamed it as the “10 images” condition to remove any assumptions of recent experience effects. The second condition (the “2 images” condition) contained two images, one of which appeared on 10% of trials, the other which appeared on 90% of trials. Therefore, we had two “main samples”. Main sample 1 was the image that most closely matched our initial task design: it had a high probability of appearing in the 2 images condition (90%) and a lower probability of appearing in the many images condition (10%). Main sample 2 was the image that had equal probability of appearing in the two conditions. Behavioral performance was calculated separately for each image in each condition, collapsed across blocks of the same condition. We report the change in *d’* as the performance in the 2 image block minus the performance in the many images block.

**Figure 3.**
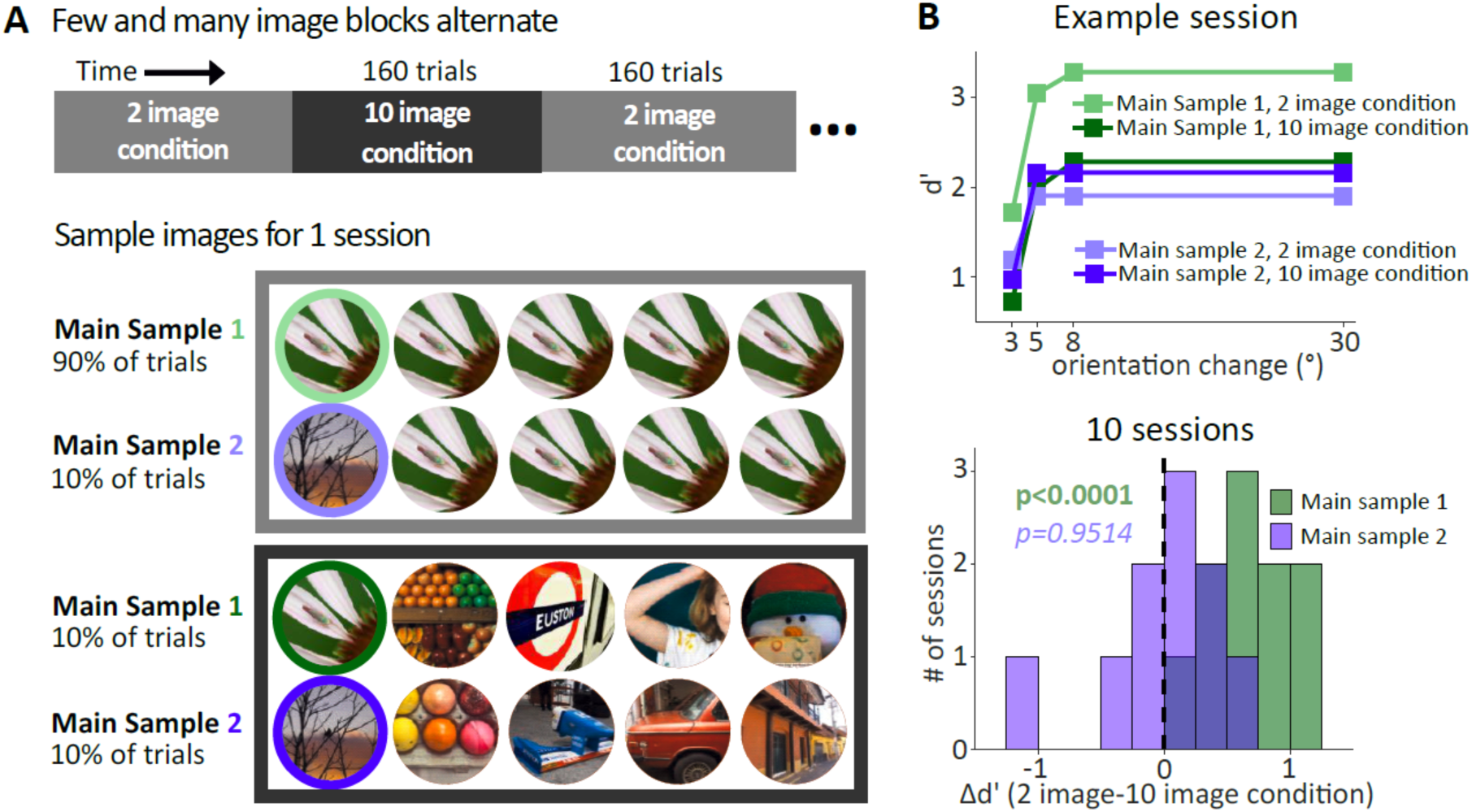
Improvement in behavioral performance for the main sample is due to stimulus probability, not the number of images in each condition. A) Modified task design to test the impact of stimulus probability vs. perceptual load. Blocks of trials with 2 possible sample images and 10 possible sample images alternated throughout the session. The 2 image condition contained two potential sample images, one of which was shown on 90% of trials (main sample 1, green), the other was shown on 10% of trials (main sample 2, purple). The 10 image condition contained 10 potential sample images each shown on 10% of trials (equivalent to original task, Fig 1). Therefore, there were two main samples. B) Behavioral results for main samples 1 and 2. Main sample 1 showed a strong effect of recent experience, but main sample 2 elicited similar behavior between blocks (top, *d’* for each main sample in the two conditions in one example session). Across 10 sessions from monkey RA (bottom), *d’* remained improved between conditions for main sample 1 but not main sample 2 (paired t-test p-values shown).

### Analysis of neuronal responses

We excluded from further analysis all neurons with a firing rate less than 1 spike per second (sp/s) to the main sample. Peri-stimulus time histograms (PSTHs) were created by aligning spike trains to stimulus onset and averaging the spike rate in 5ms time bins across trials. Because we detected effects during both the transient and sustained portion of the response (Fig 4), all analyses of neuronal firing rates used the 45-300ms window following stimulus onset. Only complete presentations of the first flash were included in the analyses for Figures 4-7. Figure 8 included all complete flashes of the main sample because the greater trial count provided by using all flashes is important given trial limitations of factor analysis (Williamson et al., 2016). Unless otherwise noted, firing rates were averaged across blocks of the same condition.

**Figure 4.**
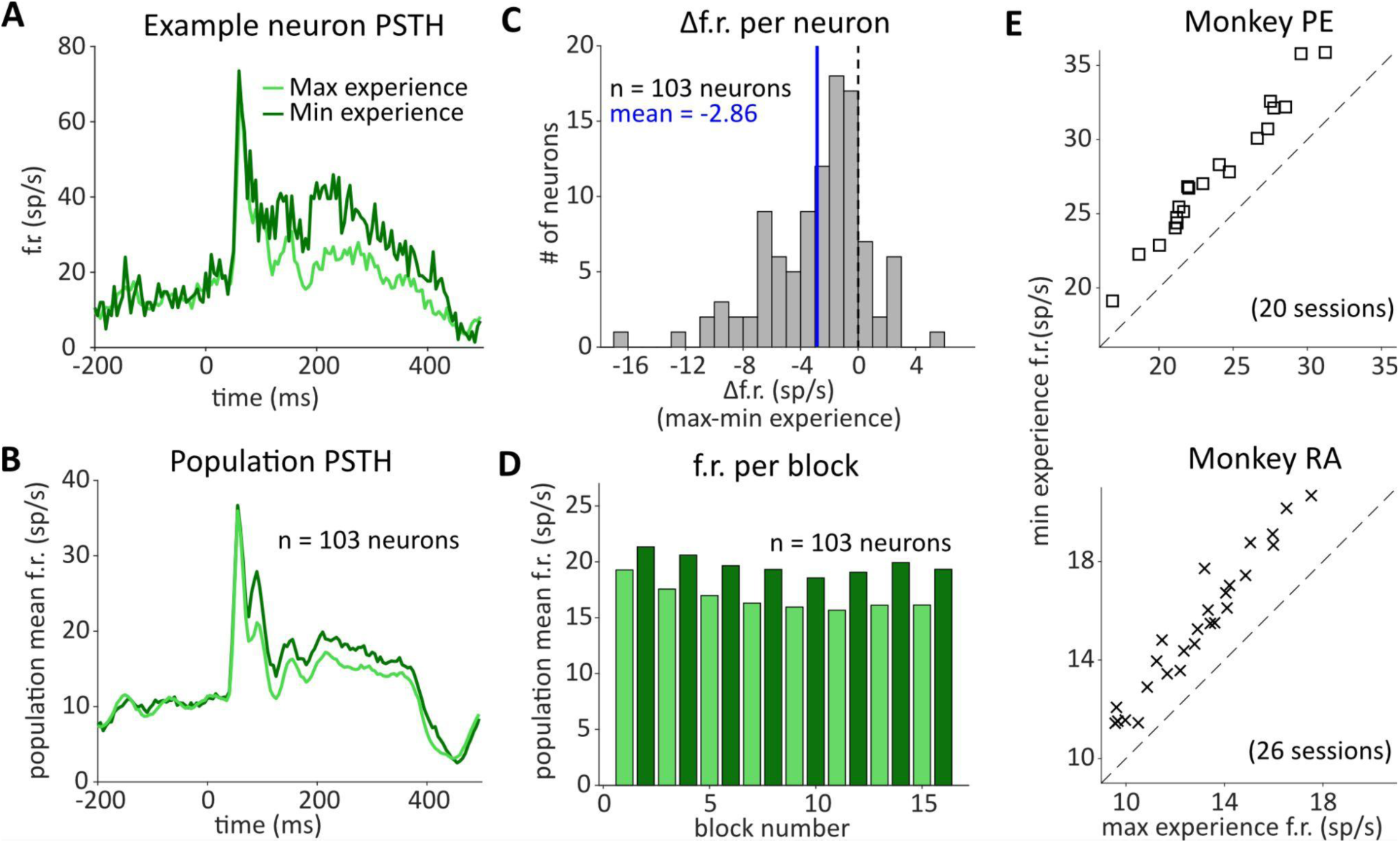
Maximizing recent visual experience is associated with decreased firing rates. A-D are from one example session from Monkey RA. Only responses to the first flash on trials of the main sample image were included. Unless otherwise noted, responses were averaged across blocks of the same condition. A) Example neuron peri-stimulus time histogram (PSTH) for the main sample image during max and min experience conditions, aligned to stimulus onset (time 0). Stimulus offset was at 300ms. The vertical axis is the firing rate (f.r.) in spikes per second (sp/s). B) Population average PSTHs across all neurons in 1 session. C) Difference in firing rate (average over an epoch 45-300ms following stimulus onset) for each neuron in an example session. Values to the left of 0 indicate that the firing rate was lower in the max experience condition. The blue line indicates the mean across neurons. D) The average firing rate across neurons for each block in an example session. E) The average firing rates for the main sample image across neurons were lower in the max than the min experience conditions for all sessions from two monkeys (p<0.0001). Each point represents one session. The dashed line indicates unity slope.

Firing rates to the target were calculated during the 45 -100ms bin following target onset on correct trials. This was to avoid any possible artifacts due to eye movements (e.g. the saccade to the target to receive a reward) which can occur after 100ms. In Figure 5B, target responses were averaged across all target difficulties.

**Figure 5.**
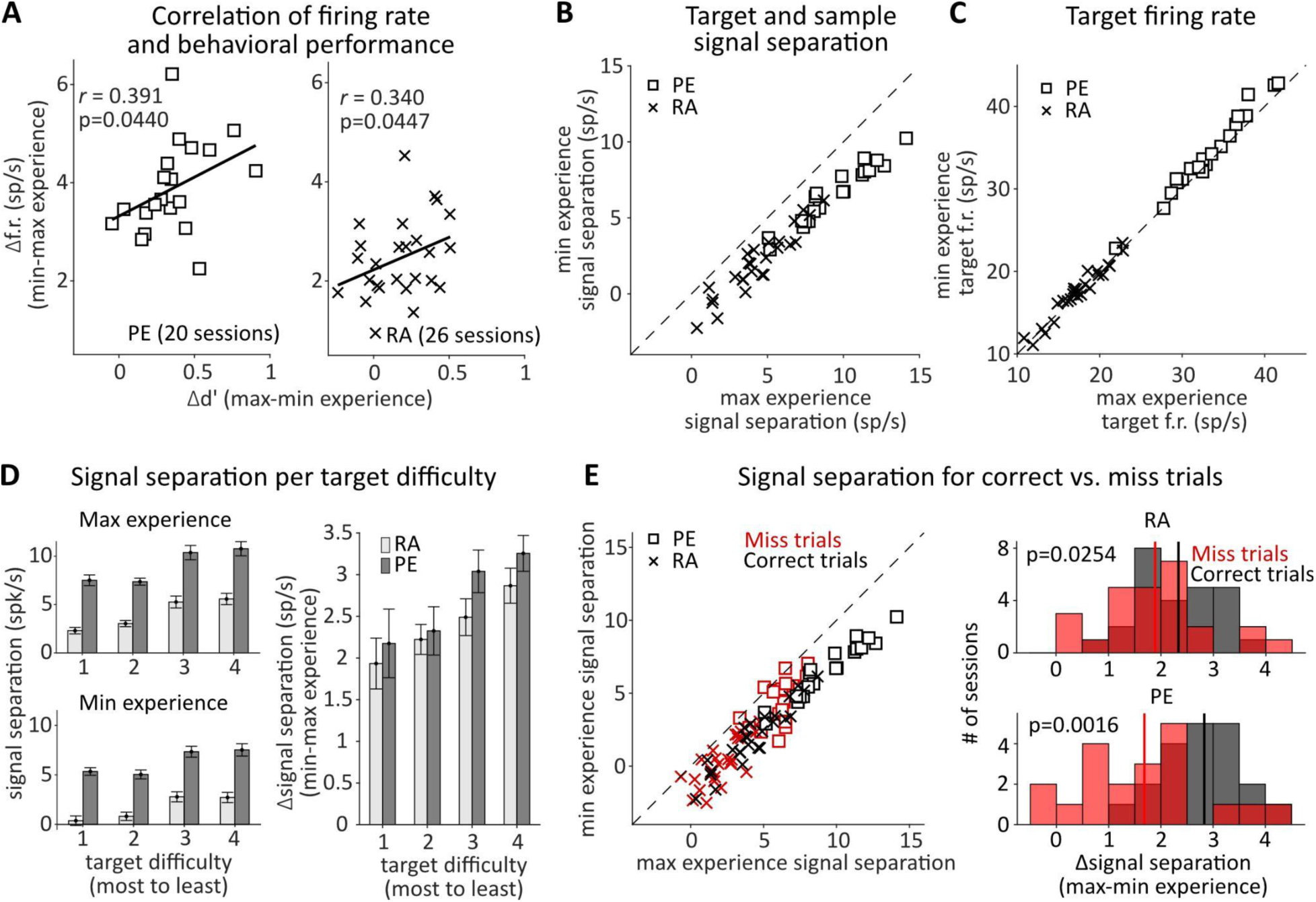
Reduction in neuronal activity is correlated with behavioral improvement and leads to larger signal separation. A) The magnitude of the firing rate difference between max and min experience was correlated with the behavioral effect in *d’* (right-tailed Pearson’s correlation). A best-fit line to the data is shown in black. The sign of the difference was chosen such that each value tended to be positive. Each point represents one session for monkey PE (left) and RA (right), all 46 sessions are shown in this and subsequent panels in this figure. B) Signal separation was calculated as the average response to the target (collapsed across the target difficulties) minus the average response to the first flash of the main sample image. Unity lines are dotted for reference. The max experience condition had a significantly higher signal separation (paired t-test p<0.0001). C) The average firing rates across neurons in each session for the target image on trials of the main sample. Each point represents one session. Target firing rates were similar in the max and min experience conditions. While the small difference was significant (paired t-test, p<0.0001, data combined across animals), this effect was present in only one animal (RA, p=0.3965; PE, p<0.0001). D) Average signal separation as a function of target difficulty for each monkey in each condition (left). Target 1 was the most difficult (smallest change), and target 4 was the least difficult (greatest change). (Right) The difference in signal separation between conditions as a function of target difficulty. Error bars show s.e.m. E) (Left) Two points per session indicating the average signal separation across miss (red) and correct (black) trials. (Right) The difference in signal separation for correct and miss trials for each subject. Vertical solid lines indicate means of each distribution.

### Comparison with within-trial stimulus repetition

We sought to relate the changes in firing rates we observed between the two conditions with firing rate changes due to within-trial repeated flashes of an identical visual stimulus. We defined “within-trial experience” as the change in firing rate between flash 1 and flash 2 in the min experience condition, and “across-trial experience” as the change in firing rate between flash 1 in the max experience condition and flash 1 in the min experience condition. For analyses involving within-trial experience, only trials with 2 or more flashes were included, even when determining firing rates on the first flash. For each neuron, we calculated the within- and across-trial firing rate effects. We measured the Spearman’s correlation between the % decrease in firing due to across-trial experience and the % decrease in firing rate due to within-trial experience in the min experience condition (Fig 6).

**Figure 6.**
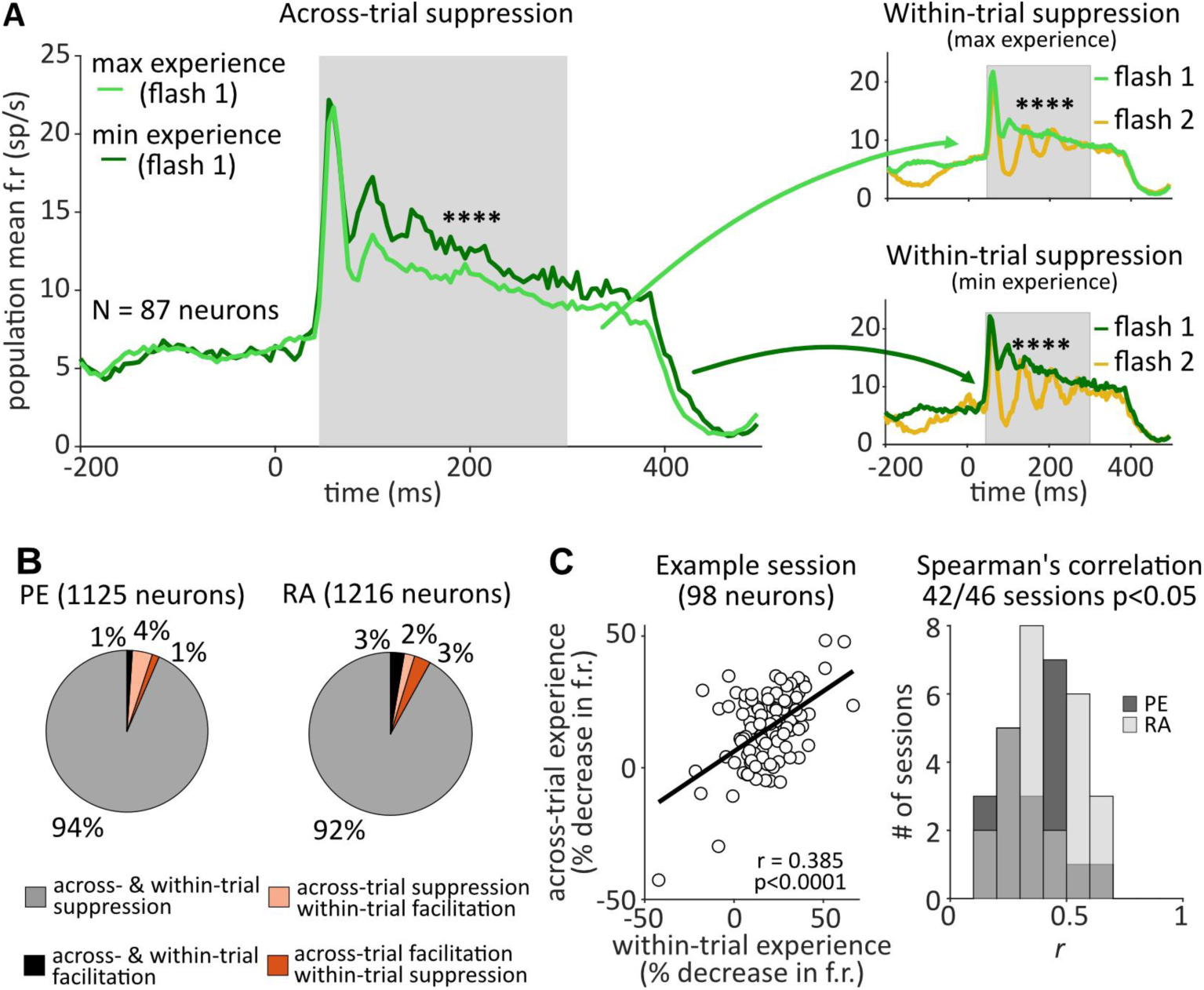
Effects of across-trial and within-trial experience are related. A) Population averaged responses on trials of the main sample image for one example session. PSTHs are aligned to stimulus onset. Left; population PSTH for the main sample in max and min experience conditions (akin to Fig 3B, but a different session is shown here) showing across-trial suppression with increased experience. Responses were calculated using the first flash of the main sample in each condition. Right; population PSTH for the first and second flash in the max (top) and min (bottom) experience conditions showing within-trial repetition suppression (i.e. adaptation). The green PSTHs are identical to those on the left. The gray shaded region indicates the time period over which responses per trial were calculated. B) The percentage of neurons falling under each combination of facilitation and suppression due to within- and across-trial experience. Only neurons with significant changes in both within- and across-trial firing rates are included. Opposing effects are in orange and represented 5% of neurons in each animal. C) The change in firing rates due to within- and across-trial experience are correlated. Left; example session showing the Spearman’s correlation between the percentage change in firing rate within trials in the min experience condition vs. the percentage change in firing rate across trials for each neuron (best-fit line in black). Right; The correlation values for all sessions for each monkey show a strong positive trend.

### Analysis of noise correlation

The pairwise noise correlation (also referred to as spike-count correlation, or r_sc_) for the main sample was calculated by taking the Pearson’s correlation between two neuron’s spike counts on each trial of the main sample. This was then repeated for all pairs of neurons, excluding pairs that were on the same array channel. Given that the number of trials can impact the estimate of noise correlation, we equalized the number of trials by randomly sub-selecting a number of trials from the max experience condition (which always had more trials) to equal those in the min exprience condition. To ensure that the sub-selection was representative of the noise correlation values we would have obtained if we included all trials, we repeated this sub-selection 2000 times, calculated the average r_sc_ for each sub-selection, and took the average across all 2000 sub-selections. Then, we chose the sub-selection of trials that was closest to the average r_sc_ across all sub-selections, and used those trials for further analysis. r_sc_ was calculated separately for each condition for flash 1. We report the mean and standard deviation of the distributions of noise correlations for the main sample during max and min experience conditions in Figure 7A.

**Figure 7.**
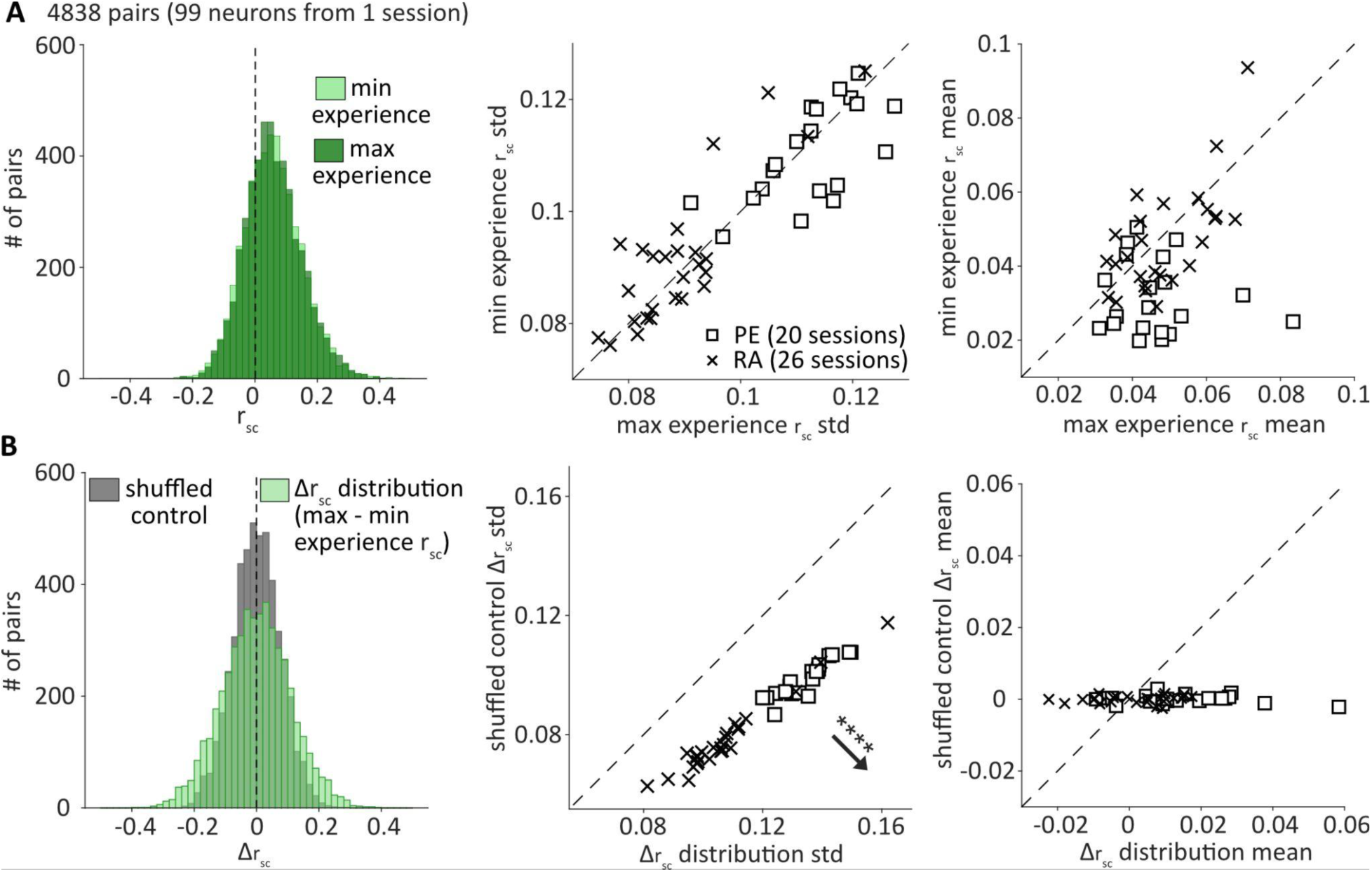
Noise correlations are modulated by recent experience. A) Left; The distribution of noise correlations (r_sc_) for an example session in max and min experience conditions. r_sc_ was calculated for the first flash of trials of the main sample. The dashed line indicates a value of 0. Middle; The standard deviation of the r_sc_ distributions were similar in both conditions across sessions for two animals. Right; The mean of the r_sc_ distributions for each session were similar in both conditions across sessions. While there was a small difference that was significant (paired t-test, p=0.0017, data combined across animals), this effect was present in only one animal (PE). B) Left; The same example session as in (A) showing the distribution of the change in r_sc_ (Δr_sc_) between max and min experience conditions for each pair (green) and the Δr_sc_ for each pair when trials were shuffled (gray). The change in r_sc_ between conditions was more variable in the real data (i.e., the distribution was wider) than in the shuffled control. Middle; The standard deviation of the Δr_sc_ and shuffled control distributions for each session for two monkeys showing a significant difference (paired t-test, p<0.0001), verifying the observation of a single session in the left panel. Right; The mean of the Δr_sc_ and shuffled control distributions for each session for two monkeys. While there was a small difference that was significant (paired t-test, p=0.0018, data combined across animals), this effect was present in only one animal (PE).

To assess if each pair’s r_sc_ value was changing between conditions, we took the difference in r_sc_ (max-min experience) and compared to a shuffled control. To obtain the shuffled control, for each neuron and each condition, we shuffled the order of trials and re-calculated r_sc_. We then took the difference in r_sc_ (max-min experience) with shuffled trials. We report the mean and standard deviation of the change in r_sc_ distribution and shuffled control in Figure 7B.

### Factor analysis

We used the dimensionality reduction method factor analysis (FA) (Churchland et al., 2010; Harvey et al., 2012; Cunningham and Yu, 2014; Williamson et al., 2016; Bittner et al., 2017; Athalye et al., 2018; Huang et al., 2019; Umakantha et al., 2021) to characterize the covariability of our neuronal population under different task conditions. FA allowed us to separate the variance shared among a population of neurons from each neuron’s independent variance, making it particularly well suited for analyzing population activity.

We performed FA on the spike of every flash of the main sample separately in max and min experience conditions. We used the spike counts during the 45-300ms time window following stimulus onset for each complete stimulus presentation (i.e. all flashes) in each trial. Only one neuron per channel was included. FA results are dependent on the number of trials, so we matched trial numbers across conditions. For each session, we sub-selected trials from the max experience condition (which has more repetitions of the main sample than the min experience condition). To ensure our results were robust to the variability that comes with randomly selecting a particular set of trials, in the max experience condition we first ran FA using 30 groups of randomly selected trials. We then calculated the mean %sv and loading similarity across groups, and identified the group that led to the smallest difference from those means. We then used this group for further analysis. For Figure 8E, we ran FA on the spike counts of every flash of the main sample separately for correct and incorrect (misses and false alarm) trials regardless of experience condition. All subsequent procedures are as described above.

**Figure 8.**
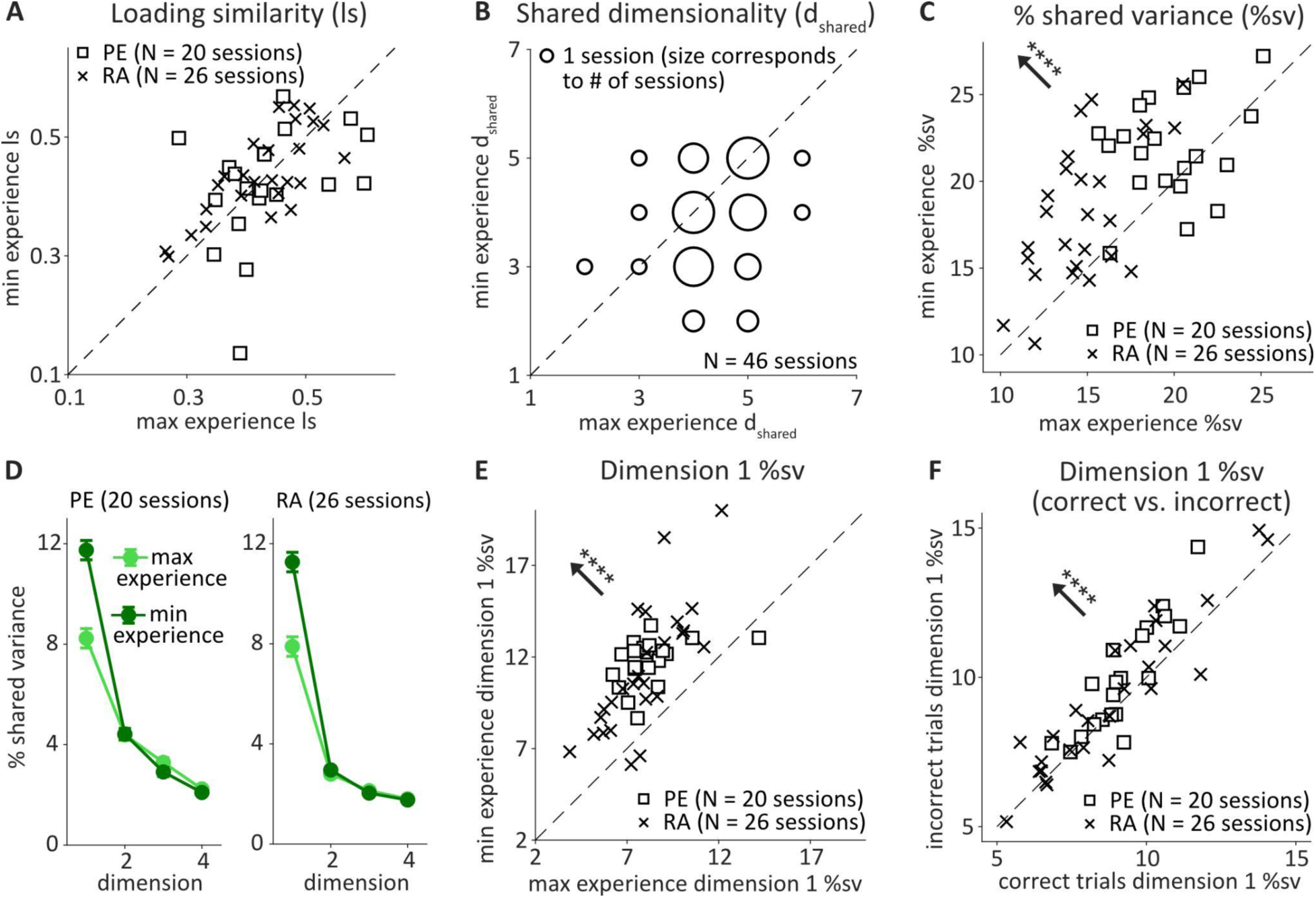
Greater recent experience is associated with a decrease in shared variability. A) Loading similarity for the first dimension in max and min experience conditions for two monkeys. Each point represents one session. B) Shared dimensionality (d_shared_) in each condition for two monkeys. The size of the circle is proportional to the # of sessions that fell at the coordinate of each plotted point (collapsed across monkeys). C) Average percent shared variance (%sv) across neurons in each condition for two monkeys. Each point represents one session. There was a significant decrease in %sv in the max experience condition (paired t-test, p<0.0001). D) %sv for each dimension following singular value decomposition. Dimensions after 4 are not plotted as there were very few sessions with a dimensionality greater than 4. For each x-axis value, only sessions which had an identified dimension of that number contributed to the data shown. The points indicate the mean and standard error across sessions. E) The average %sv across neurons for the first dimension was significantly lower in the max experience condition (paired t-test, p<0.0001). Each point represents one session. F) The %sv for the first dimension on correct vs. incorrect trials showed that correct trials had a significantly lower %sv (paired t-test, p=0.0001).

FA is defined as:

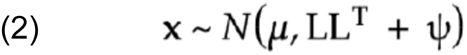

Where **x** is a *n* x 1 vector of spike counts with n as the number of neurons, **μ** is a *n* x 1 vector of mean spike counts. *LL^T^* is the shared component, where *L* is a *n* x *i* loading matrix with *i* as the number of latent dimensions, and Ψ is a *n* x *n* diagonal matrix containing each neuron’s individual variance. To determine the number of latent dimensions *i*, we ran FA using different numbers of latents and chose the value of *i* that maximized the cross-validated log-likelihood.

% shared variance refers to the percentage of the total variance that is shared versus independent (Williamson et al., 2016). The % shared variance for neuron n is calculated as:

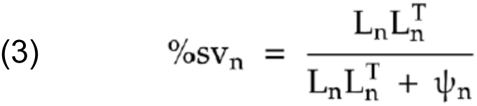

Where L_n_ is the loading matrix for each neuron, and Ψ_n_ is each neuron’s individual variance.

In figure 8C we reported the average %sv across neurons. In figure 8C-E we used singular value decomposition to partition the shared variance along each latent. We then computed the %sv for each neuron *n* along each latent *i* as:

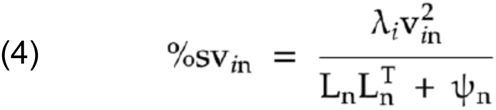

Where λ is the eigenvalue of *LL^T^* corresponding to the *i*th latent, and *v*_in_ is the *n*th entry in the *i*th eigenvector of *LL^T^*. We then averaged this value across all neurons to determine the %sv along each dimension. In figure 8D-E we report the average %sv across neurons for the first dimension.

Loading similarity is the similarity of loading weights across neurons for each latent (Cowley et al., 2020). We compute loading similarity as:

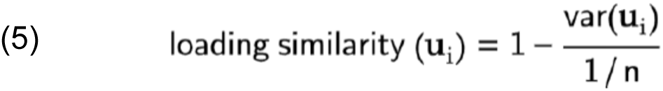

Where *n* is the number of neurons, and u_i_ is an *n* x 1 vector of weights for a given latent *i*. A value of 1 indicates that the weights for all neurons are the same (either all positive or all negative). A value of 0 indicates that the loading weights are as variable as possible. We report the loading similarity for the first latent (the dominant dimension).

Shared dimensionality (referred to as simply “dimensionality” in the results, or d_shared_) was computed as the number of latents needed to explain 95% of the shared covariance, *LL^T^* (Williamson et al., 2016).

### Firing rate controls for shared variance

Firing rates can affect %sv, with higher firing rates leading to lower %sv values. Therefore, we sought to control for population firing rate differences between the max and min experience conditions by sub-selecting neurons in each condition that would result in matched population firing rates. From the analyses described above, we obtained %sv values for each individual neuron in each condition. We also calculated the firing rate for each neuron and ranked them from lowest to highest, separately for each condition. We removed one neuron at a time from each condition, recalculated the difference in population firing rate, and repeated until the difference was reduced while keeping as many neurons as possible. The identity of the neurons in each condition may have differed, but the total number of neurons were matched. Using this method, we significantly brought down the difference in firing rate between max and min experience conditions in each session (difference in firing rate was <1 sp/s for each session). Then, we re-calculated the average 1st dimension %sv for each condition. We found in these firing rate matched controls that the max experience condition still had a lower %sv as in the unmatched data, and the results for Fig 8C-D followed the same overall pattern. We repeated this procedure for Figure 8E, and likewise found the results of the firing rate matched control were similar to those shown in Figure 8.

## Results

### Effects of recent experience on task performance

Two monkeys performed a natural scene change detection task under conditions in which recent visual experience with a particular image was either maximized or minimized (Fig 1). On every trial, a sample image flashed on the screen, and the monkey had to detect when a feature (either color or orientation) of the sample image had changed. There was a fixed probability of a change happening on every flash (34% or 36%, see *Methods*), meaning that ∼35% of trials consisted of one flash of the sample image and then the target, ∼23% of trials had two flashes of the sample image then the target, and so on. Because there was a fixed probability of a change, the animal could not anticipate when the change would occur. In the condition with maximal recent experience (hereafter referred to as “max experience”), a single image (the main sample) was used on every trial. In the condition with minimal recent experience (hereafter referred to as “min experience”), the main sample was used on only 10% of trials (with 9 other images comprising the other 90% of trials). By using recent experience to modulate the probability of the main sample being the sample image on a given trial, we also modulated the animal’s expectation of an upcoming image. The majority of trials were short, with more than 70% having 3 or fewer sample flashes (sample plus interstimulus time spanning 0.5 to 1.5 seconds). Thus a decision of whether the image changed or not had to be made quickly, and an expectation of which image would be presented on a given trial could confer a behavioral advantage. Each day, a new main sample and accompanying 9 images were chosen (see *Methods* for a description of this process), and a change type was selected (either color or orientation). Therefore, our results generalize across different images and different feature changes, and could not occur due to across-day perceptual learning. There were 4 change difficulties within each type (e.g., 4 different magnitudes of orientation change), and max and min experience blocks of trials alternated throughout the day. Additionally, we alternated which block type (i.e. max or min experience) was the starting block for the day.

To determine if greater recent experience improves the ability to detect a change in an image, we analyzed behavior in terms of discriminability (d-prime, or *d’*), criterion, and reaction time. Importantly, we compared the performance to the main sample image in max and min experience conditions, not to all images in the two conditions. Both monkeys showed a robust increase in *d’* for the main sample image when maximizing recent experience, with 77% of sessions (20/26) for monkey RA and 95% of sessions (19/20) for monkey PE having better performance in the max experience condition (Fig 2A-B). We did not see a consistent effect of recent experience across animals for criterion (Fig 2C; paired t-test; monkey RA showed no difference, p=0.4086; monkey PE showed an increase in criterion, p<0.0001), nor reaction time (Fig 2F; paired t-test; monkey RA showed a small decrease in the max experience condition, p=0.0340; monkey PE showed no difference, p=0.3393), indicating that the predominant effect of experience was a change in sensitivity or *d’*. The value of *d’* is calculated from hit rate and false alarm rate (Fig 2D-E), and *d’* can increase through a mixture of those two values. For example, an increase in hit rate while maintaining a constant false alarm rate, as in monkey RA, or a decrease in false alarm rate while maintaining a constant hit rate, as in monkey PE, both indicate an improvement in sensitivity to the changes in the stimulus. Taken together, the changes in *d’* were the most consistent effects between animals and our results show that in situations where subjects have extensive recent experience with an image, they demonstrate improved abilities to detect changes in that image.

### Alternative explanations for behavioral effects of recent visual experience

We refer to our manipulation as a modulation of recent experience because the probability of the main sample appearing on each trial can be determined based on the recent trial history. We found that overall behavioral performance was better in blocks where one image was shown on every trial (100% probability) as opposed to on fewer trials (10% probability). However, there are two potential alternative explanations of this behavioral result, which we addressed with additional analyses and experiments.

First, an increase in task difficulty can improve an animal’s performance in a change detection task - an effect that has been attributed to an increase in effort or motivation (Spitzer et al., 1988; Boudreau et al., 2006; Ruff and Cohen, 2014b; Ghosh and Maunsell, 2021). In the min experience blocks containing ten different images, new images were selected each day without measuring behavioral thresholds in advance. Because of this, the difficulty of the ten image block varied from day to day based on the difficulty of the change detection task with each of the ten natural images. To consider the role of task difficulty, we assessed the difficulty of the main sample *relative* to the other images used that day. In some sessions the main sample had better change detection performance than the other images (30/46 sessions) and in other sessions the opposite was true (16/46 sessions). On days where the main sample was particularly challenging for change detection, the performance with that main sample had some tendency to be better in the max experience condition in one animal (Pearson’s correlation, PE *r*=0.458, p=0.0186; RA *r*=0.284, p=0.2250). This indicates that the difficulty of the 10 image condition may have impacted the day-to-day effects of experience. However, our study focused on a matched comparison of performance for the main sample within each session, and we found that performance for the main sample was still better when it was experienced on every trial (39/46 sessions) irrespective of the day-to-day changes in difficulty (in 33/46 sessions the main sample was more difficult than other images, and in 13/46 sessions the main sample was less difficult than other images). These results suggest that although difficulty varied day-to-day, a difference in difficulty was not the main driver of the improvement in discriminability for the main sample in the max experience condition.

A second explanation for the behavioral performance difference in the max and min experience conditions is related to perceptual load and working memory. Although our intention was to focus on differences in recent experience induced by the two conditions, it is possible that the two conditions differed in how working memory was used to store information about the images presented in each trial. Perhaps the difference in performance on the main sample was not because the animal had a perceptual improvement in the max experience block, but rather that there was a perceptual deficit induced by the presence of the additional images in the min experience block. To determine if this could be the case, we designed a control experiment that would allow us to distinguish between the effects of varying recent stimulus experience and varying the number of images in each condition (which could create differences in working memory, perceptual load, etc.). This involved a small modification to our original task design (Fig 3A). The 10 image condition remained as is, with 10 different images each being shown on 10% of trials. Instead of a 1 image condition, we used a 2 image condition, where one image (main sample 1) was used on 90% of trials, and a second image (main sample 2) was used on 10% of trials. Both main samples were also found in the 10 image condition.

In this control experiment, the stimulus probability of main sample 1 is akin to our original experiment, with only a decrease from 100% to 90% in the 2 image condition, and a 10% probability in both experiments in the 10 image condition. Main sample 2, however, had equal stimulus probability in both conditions (10%). For main sample 1, we recapitulated our initial finding: there was a significant improvement in behavioral performance for main sample 1 in the 2 image (higher stimulus probability) condition compared to the 10 image condition (Fig 3B, paired t-test, p<0.0001). This indicates that the reduction from 100% to 90% stimulus probability in this control experiment did not remove the effect of maximizing recent experience. The key additional comparison this control allowed us to perform was in the behavior of the animal for main sample 2 in the two conditions. The stimulus probability of this image was equal in both blocks, but the perceptual load from working memory demands might be considered higher in the 10 image condition. However, there was no significant difference in behavioral performance for main sample 2 across these conditions (Fig 3B, paired t-test, p=0.9514). These results support our initial interpretation: the improvement in discriminability of the main sample in the 1 image maximal experience condition was not due to an impairment in performance when a large number of images were used, but rather due to the greater amount of recent experience created through increased stimulus probability.

### Effects of recent visual experience on firing rates

Single neuron firing rates in the visual system reflect both the preferences of each neuron for visual input as well as modulation by numerous cognitive and contextual factors. We sought to understand the way in which recent experience of a particular visual input modulates firing rates of individual neurons. There are two timescales of recent visual experience in our study: across-trial accumulation of experience (the key difference between the min and max experience conditions) and within-trial repetition of the visual input (typically studied under the umbrella of adaptation). Both will be addressed, although the focus of this study is on the former since the behavioral effect we are studying is across trials within a session. To isolate our analysis from well-known effects of repeated presentations of a stimulus over tens to hundreds of milliseconds (typically labeled as short-term adaptation or repetition suppression), we focused on the responses to the first visual stimulus presentation on each trial (referred to as the first flash). The first flash is the time point where the two conditions differed the most in recent experience, and therefore most likely to be related to behavioral differences between the two conditions.

When we compared the firing rates for the first flash of the main sample (45-300ms after image onset) in max and min experience conditions we found, in line with past studies of expectation, that average firing rates were decreased with max experience (Fig 4). The majority of neurons exhibited suppression of varying magnitudes, although some neurons had enhanced firing rates in the max experience condition (Fig 4C). The decrease in firing rate was robust throughout the session as we alternated blocks of each experience condition (Fig 4D), and this effect was evident in every session in both monkeys (Fig 4E). Thus, the predominant effect of maximizing recent experience on the firing rates of individual neurons was one of suppression.

We next sought to link these firing rate changes to the behavioral effects of experience that we observed in each animal. We analyzed this on a session-by-session level, measuring the firing rate difference along with the difference in behavioral *d’*. We found that these two measures were significantly correlated across sessions in both monkeys (Fig 5A; right-tailed Pearson’s correlation; Monkey PE r=0.391, p=0.0440; Monkey RA r=0.340, p=0.0447), indicating that in sessions with a larger suppression of firing rates, there was also a greater improvement in discriminability of the main sample image. Thus, suppression of firing in individual neurons may represent a neural correlate of recent visual experience.

Next, we looked to identify a possible mechanistic explanation for how suppression of firing rates could lead to improved stimulus discriminability. Theoretical frameworks such as predictive coding have suggested that suppression of activity to an image could make a new incoming image more salient, as the difference in the signal would be greater and therefore more easily decoded by a higher cortical area (Walsh et al., 2020). While we found a clear impact of experience on the neuronal response to the main sample image, the ability of the visual system to discriminate between two images should also depend on the response to the target (i.e., the main sample image after the change in color or orientation). We therefore considered whether the changes in response to the main sample led to a greater separation between responses to sample and target. To do this, we subtracted the population average response to flash 1 of the main sample from the response to the target (collapsed across all target difficulties) in each condition. We found a robust increase in the separation between sample and target in the max experience condition (paired t-test, p<0.0001, Fig 5B). We chose to use flash 1 in our analysis as opposed to the flash preceding the target as flash 1 is the point in which the difference in experience is greatest between conditions. When we repeated the analysis using only trials with 1 main sample flash (where by definition the first flash was the flash preceding the target) we obtained the same result (paired t-test; RA p<0.0001; PE p=0.0017). The increase in signal separation appeared to be largely driven by the modulation in response to the main sample, because firing rates to the target stimulus were overall very similar in the two conditions (Fig 5C, although there was a small effect on target response in one animal - paired t-test; Monkey PE, p<0.0001; Monkey RA, p=0.3965).

We then looked at the signal separation per target difficulty and found that, as expected, in each condition the difference between sample and target response was larger for more easily detectable targets (Fig 5D, left) which represented more substantial changes from the main sample image. In addition, the finding that signal separation was greater in the max experience condition was robust across target difficulties (Fig 5D, right). Next, to determine if signal separation was reflective of behavior, we examined correct and miss trials separately. We found that in both conditions correct trials had greater signal separation than miss trials (Fig 5E, left, black points had higher values than red points), and, the difference in signal separation between min and max experience conditions was significantly greater on correct trials (Fig 5E, right, paired t-test RA p=0.0254, PE p=0.0016). This indicates that when recent experience had a larger effect of increasing signal separation in the population of neurons, the animal was more able to correctly detect a change in the image. Together, these results indicate that there was a robust increase in signal separation between the main sample image and the target images - an important potential neural substrate for the behavioral effects of experience - which was largely driven by changes in the neuronal responses before the target image was shown.

### Interactions between within and across-trial experience

One possible mechanism underlying the decrease in firing rates we observe with heightened experience is that of short-term adaptation. Adaptation refers to the changes in neural activity that occur when a sensory input is repeated (Solomon and Kohn, 2014; Webster, 2015). Suppression of neural responses, i.e. repetition suppression (Grill-Spector et al., 2006; Auksztulewicz and Friston, 2016), is most common and can occur at various timescales ranging from several hundreds of milliseconds to many seconds following prolonged exposure. Although our task design involved eye movements between trials that provided variety to the visual inputs, and there were variable lengths of pauses between trials depending on how quickly the animal engaged in the task, it is still the case that in the max experience block the recent visual input was dominated by repeated presentations of the main sample image. Thus, the well-studied effects of short-timescale adaptation on neurons could be the means by which across-trial recent experience modulated firing rates and in turn behavior. If the firing rate changes seen between max and min experience conditions were related to within-trial adaptation, we would make the following predictions: 1) for each neuron, the sign of firing rate change due to across-trial and within-trial experience should match (e.g. a neuron whose firing rate decreases with within-trial adaptation should also have a firing rate decrease across trials), 2) the amount of firing rate change due to across-trial and within-trial experience should be correlated across neurons.

To assess across-trial effects, we focused our analysis on the difference in firing rates between flash 1 of the main sample in the two conditions (as in Fig 4 & 5). We refer to this difference as “across-trial” experience (Fig 6A, left). Constraining to just the first flash allowed us to better isolate the effect of experience prior to any short-term (within-trial) adaptation effects. To assess the effect of short-term adaptation, we focused our analysis on the difference in firing rates between flash 1 and flash 2 of the main sample within a trial in each condition (Fig 6A, right). We refer to this difference as “within-trial” experience which can take the form of repetition suppression or repetition facilitation. When comparing the effects of within- and across-trial experience (Fig 6B & 6C), we used each neuron’s within-trial adaptation value from the min experience condition, as the adaptation found in that condition was expected to be solely confined to the time scale of individual trials since the sample images varied from trial to trial.

First, if across-trial effects were the result of accumulated within-trial adaptation, we would predict that neurons exhibiting within-trial repetition suppression would exhibit across-trial suppression, and neurons exhibiting within-trial facilitation would exhibit across-trial facilitation. For each neuron, we assessed whether firing rates were increased or decreased with within-trial and across-trial experience. We then only considered neurons that had significant (t-test p<0.05) changes in activity both within and across trials. The vast majority of neurons (94% for PE, 92% for RA) exhibited within- and across-trial suppression (Fig 6B), although there were a small minority of neurons that did not show the same sign in firing rate change within and across trials (5% for PE, 5% for RA). This result indicates that suppression is the most prevalent firing rate change for both within-trial and across-trial recent experience. Second, we looked to see if the magnitude of firing rate changes due to within-trial and across-trial experience were correlated. For each session, we measured the correlation between each neuron’s within-trial effect (min experience flash 1 firing rate - min experience flash 2 firing rate) and across-trial effect (min experience flash 1 firing rate - max experience flash 1 firing rate). For the majority of sessions, within- and across-trial experience were correlated across our populations of neurons (Fig 6C; 96% of sessions Monkey RA, 85% of sessions Monkey PE, Spearman’s correlation p<0.05), indicating that neurons that have larger within-trial changes in firing rates also tended to have larger across-trial changes in firing rates. Together, these results suggest that short-term adaptation mechanisms may accumulate to produce longer across-trial changes in neural activity and in turn impact behavioral performance.

### Effects of recent experience on neuronal populations

Our analysis so far has focused on changes in firing rates of individual neurons and the population average. However, in addition to changes in firing rates, experience could be acting on population activity to change the structure of variability shared among groups of neurons. Studies of other cognitive factors such as learning (Gu et al., 2011; Jeanne et al., 2013; Ni et al., 2018), attention (Cohen and Maunsell, 2009; Mitchell et al., 2009; Herrero et al., 2013; Snyder et al., 2016), and task context (Cohen and Newsome, 2008; Bondy et al., 2018) report a reduction in pairwise noise correlations (the trial-by-trial correlations between the responses of pairs of neurons) is associated with improved perceptual abilities. Therefore, to investigate if changes in recent experience affect noise correlations (also known as spike count correlations, or r_sc_), we first compared the mean and standard deviation of the distributions of r_sc_ for the main sample during max and min experience conditions. On average, noise correlations were slightly positive in both max and min experience conditions (an average of 0.048 and 0.040, respectively), consistent with previous work showing small positive values for mean r_sc_ (Cohen and Kohn, 2011). However, when comparing the max and min experience conditions, we found no difference across all pairs of neurons in the standard deviation of the r_sc_ distribution (Fig 7A, middle), and an inconsistent effect on mean r_sc_ (Fig 7A, right; paired t-test; Monkey RA no difference in r_sc_ mean, p=0.4427; Monkey PE a small increase in r_sc_ mean, p=0.0007). Therefore, in contrast to studies of other cognitive factors, we do not find evidence that recent visual experience alters the mean noise correlation or shifts the standard deviation.

Even though the overall distribution of noise correlations did not appear to be affected by recent experience, it is possible that correlations between individual pairs were still changing between conditions in such a way as to maintain the overall distribution of r_sc_. This would represent changes in the neuronal population variability that might remain hidden from summary statistics of r_sc_ distributions (Umakantha et al., 2021). We sought to determine if r_sc_ for each pair of neurons was changing between conditions. For each pair, we compared the difference in r_sc_ between conditions (max-min experience r_sc_) to a shuffled control where we shuffled the trial order for each neuron in each pair, recomputed r_sc_ for both the max and min experience conditions and took the difference (max-min experience of shuffled data). We did not see consistent effects on r_sc_ mean (paired t-test; Monkey RA p=0.4411; Monkey PE p=0.0009), but in every session we found an increase in the standard deviation of the difference in r_sc_ distribution compared to the shuffled control distribution (Fig 7B). This indicates that more pairs of neurons exhibited changes in r_sc_ between conditions in the real data than would be expected by chance in the shuffled control. Together, these results indicate that co-fluctuations between neurons are indeed changing between max and min experience conditions, but the changes are not well captured by a difference in the mean and standard deviation of pairwise noise correlations across conditions.

Because population-level changes in variability can be sometimes difficult to detect in distributions of r_sc_ (Umakantha et al., 2021), we considered an alternative approach – factor analysis (FA) – that would allow us to characterize the structure of population covariability (Cunningham and Yu, 2014). FA is a dimensionality reduction method that has revealed changes in population covariability during decision making (Harvey et al., 2012; Mante et al., 2013; Kaufman et al., 2015), learning (Sadtler et al., 2014; Ni et al., 2018; Vyas et al., 2018), and attention (Cohen and Maunsell, 2010; Rabinowitz et al., 2015; Snyder et al., 2018; Huang et al., 2019; Umakantha et al., 2021). We analyzed the shared variability of neuronal populations separately in the two conditions, max and min experience, using factor analysis applied to the responses gathered from every flash of the main sample (see *Methods*). To characterize the structure of population activity, we looked at 3 commonly reported metrics from FA: percent shared variance, shared dimensionality, and loading similarity. Percent shared variance (%sv) is a measure of the strength of shared variability and was calculated as the average %sv across all neurons in each session. Shared dimensionality (d_shared_) is a measure of the number of activity patterns present in the population activity and was calculated as the number of dimensions needed to explain 95% of the shared variance. Loading similarity (ls) is a measure of the degree to which neurons co-fluctuate together and was calculated for the first dimension.

There was no significant difference in loading similarity (Fig 8A, paired t-test; p=0.9144), and only a small difference in shared dimensionality which was driven by subject PE (Fig 8B, paired t-test; together p=0.0022, RA p=0.1613, PE p=0.0031). The most consistent result we observed was a decrease in %sv across both animals in the max experience condition (Fig 8C, paired t-test, together p<0.0001, PE p=0.0153, RA p<0.0001). Percent shared variance is calculated as the percent of variance shared across neurons divided by the sum of shared variance and individual variance. We also employed a mean-matching procedure to insure that firing rate changes did not impact this result (see *Methods*), and found it was robust to this control. The average private variance (i.e., individual to each neuron) was not different between max and min conditions in this mean-matched control analysis (paired t-test, RA p=0.6762; PE p=0.7095), indicating the main effect we observed was in the structure of shared population variability.

Recent studies have indicated that variability in cortex is primarily low-rank (i.e., a variability structure in which most of the variability is confined to one dimension). A reduction in low rank variability has been associated with a variety of sensory and cognitive factors that are linked to improved perception (Ni et al., 2018; Huang et al., 2019; Ruff et al., 2020; Umakantha et al., 2021). To determine if recent experience likewise modulates low-rank variability, we used singular value decomposition to assess the %sv along each dimension (Fig 8D). In both monkeys, we found a robust and significant decrease in %sv along the first dimension in the max experience condition (Fig 8E; paired t-test, p<0.0001), indicating that greater recent experience is associated with a decrease in low-rank variability. Additionally, we wondered if a decrease in low-rank %sv was more generally related to improved behavioral performance. We ran FA separately on correct trials and incorrect (misses and false alarm) trials (see *Methods*) and found that low-rank %sv was lower on correct trials than incorrect trials (Fig 8F; subject combined paired t-test p=0.0001; PE p=0.0035; RA p=0.0188), indicating that a decrease in shared variability is associated with correct performance on our perceptual task within the trials of each session. Together with existing literature on changes in variability due to other cognitive factors such as attention, this finding supports the notion of shifts in low-rank variability as a canonical mechanism by which reshaping cortical activity can impact behavior.

## Discussion

In this study, we sought to identify and link neural signatures to the behavioral improvements associated with recent visual experience. We trained monkeys to perform a natural image change detection task where we modulated the probability of encountering a particular image to create different levels of visual experience that would impact behavioral performance. At the neuronal level, we found that greater recent experience was associated with a suppression of neuronal activity that increased signal separation and was correlated with improved task performance. The suppression of activity across trials shared similarities to within-trial repetition effects. At the population level, greater recent experience was associated with modulation of noise correlations and a decrease in shared variability. Together, these neural effects were linked with the behavioral improvements associated with recent visual experience.

In our experiment, different levels of visual experience led to not just perceptual shifts, but a measurable improvement in perceptual abilities. Critically, this allowed us to relate neural findings to a concrete behavioral outcome. At the same time, the introduction of a task also introduces cognitive factors that impact performance, including expectation (Summerfield and de Lange, 2014; de Lange et al., 2018; Feuerriegel et al., 2021), feature attention (Maunsell and Treue, 2006; Liu, 2019), task difficulty (Spitzer et al., 1988; Boudreau et al., 2006; Ruff and Cohen, 2014b), working memory (Myers et al., 2017), perceptual learning (Gilbert et al., 2001; Tsodyks and Gilbert, 2004; Sagi, 2011), and familiarity (Sheinberg and Logothetis, 2001; Freedman, 2005; Mruczek and Sheinberg, 2007). Studies of perceptual learning and familiarity have similarities to our work, but the mechanisms that underlie those phenomena are unlikely to explain our behavioral effect. Perceptual learning is typically seen for a specific feature and across longer timescales from days to months (Gilbert et al., 2001; Tsodyks and Gilbert, 2004; Sagi, 2011) whereas our results are detectable between blocks of trials within a session (even the first block, e.g. Fig 4D) and were robust to entirely new natural images chosen each day without regard to specific image features. Familiarity is typically described as the total accumulated experience with a stimulus, sometimes within a session (Li et al., 1993) but largely across many sessions (Freedman, 2005; Mruczek and Sheinberg, 2007; Anderson et al., 2008; Woloszyn and Sheinberg, 2012; Meyer et al., 2014; Koyano et al., 2023), meaning that familiarity can only increase with each subsequent presentation. This means that familiarity for the main sample in our task increases throughout the entire session, and it is therefore unlikely to account for the decrease in behavioral performance in min experience blocks. In other words, if the subject had already built up familiarity with an image at the end of a max experience block, it would not suddenly become “less familiar” in the min experience block, and thus could not explain the decrease in performance. We also considered whether working memory load could be affected by our task conditions, with more images in the min experience condition leading to more demands on working memory and therefore worse performance. To address this possibility, we ran an additional experiment that allowed us to separate the effect of performing the task with many images from the frequency of repeating a particular stimulus (i.e. the amount of recent experience). Our results indicated that it is in fact higher stimulus probability, and not the lower number of images, that led to a behavioral improvement in the max experience condition, thus making an increase in recent experience the most likely explanation for the improvement in change detection.

While reports of changes in neural response and perception due to repeated sensory experience are widespread, there is limited data showing that adaptation leads to an overall improvement in behavior and perception (Kohn, 2007; but see Dragoi et al., 2002; McDermott et al., 2010; Wissig et al., 2013), and even some evidence of behavioral worsening (Jin et al., 2019). Although expectation leads to substantial improvements in perceptual behavior and also reductions in neuronal response (Summerfield and de Lange, 2014; de Lange et al., 2018), studies that include both neural and perceptual measures only rarely use tasks where the expectation is relevant to task performance (Kok et al., 2012; Bell et al., 2016). Broadly, studies of anticipatory prediction signals in early visual cortex have been inconsistent, with some evidence for anticipatory signals in V1/V2 in an orientation discrimination task (Goris et al., 2017), but a lack of temporal prediction signals in V1/V4 (Solomon et al., 2021). This latter study did identify prediction signals in EEG only when subjects were instructed to look for violations in a sequence, which may suggest that our task design in which recent experience was particularly relevant to task performance may have been critical to our pattern of results. Overall, our study is one of a very few to directly show that a reduction of neural activity in a sensory area after accumulated visual experience is associated with improved performance on a perceptual task.

We found that across trials, the behavioral improvement with greater recent experience was correlated with a decrease in population average firing rate. Given that both short-timescale adaptation due to stimulus repetition and across-trial buildup of expectation are reported to decrease neural activity, there have been numerous attempts to determine how adaptation and expectation interact with each other with inconsistent results (Feuerriegel et al., 2021). Additionally, only rarely have adaptation studies assessed if adaptation effects can extend across trials separated by eye movements (McMahon and Olson, 2007; Brunet et al., 2014). Our finding linking within-trial short-term adaptation to across-trial accumulated experience provides a potential link across time scales between adaptation and expectation. Because our max experience condition evoked an accumulated neural effect that appeared to build on short-term adaptation, we demonstrated that even briefly presented sensory experiences accumulated to produce longer term impacts on neuronal responses.

How might the decrease we observed in single neuron responses benefit sensory processing? Theoretical frameworks such as predictive coding posit that a decrease in firing rate to an expected stimulus would allow for a larger difference in activity to a new sensory input, thereby making the input more easily detectable (Auksztulewicz and Friston, 2016; Walsh et al., 2020). Such a process could be implemented by a hierarchical framework analogous to the structure of the primate visual system (Chao et al., 2018). Our results provide support for this idea. We measured the difference in response between the sample image and the target image as an index of the signal-to-noise ratio – a potential measure of detectability of the target by our neuronal population. We found that this signal separation was higher with more experience, and also higher on correct trials, indicating that it may reflect a neuronal substrate for stimulus detection that is shaped by sensory experience.

The relationship between neural activity and behavior is more complex than just changes in firing rates of individual neurons, with numerous cognitive factors affecting the structure of population activity. For example, repeated presentation of stimuli with different levels of uncertainty impacted the geometry of population representations in early visual cortex (Hénaff et al., 2020), with more natural stimulus transitions leading to different neural response trajectories than unnatural stimulus sequences (Hénaff et al., 2021). Dimensionality reduction, which allows for the characterization of several distinct features of population-wide covariability (Cunningham and Yu, 2014), has been used to investigate changes in neuronal populations during decision making (Harvey et al., 2012; Mante et al., 2013; Kaufman et al., 2015), learning (Sadtler et al., 2014; Ni et al., 2018; Vyas et al., 2018), and attention (Cohen and Maunsell, 2010; Rabinowitz et al., 2015; Snyder et al., 2018; Huang et al., 2019; Umakantha et al., 2021). We used factor analysis (a dimensionality reduction method) to interrogate the effects of recent experience on population activity and found that maximizing recent experience was associated with a decrease in shared variability, particularly along the first dimension. A reduction in % shared variance could result in a decrease in the overlap between the representation of two stimuli, which would allow more accurate decoding of responses by a subsequent brain area (Umakantha et al., 2021). This is supported by findings that low-rank decreases in shared variability are associated with multiple cognitive processes that impact behavior such as learning (Ni et al., 2018), attention (Huang et al., 2019; Umakantha et al., 2021), task belief (Xue et al., 2022) and task switching (Ruff et al., 2020). Our results provide further evidence that different cognitive factors, perhaps through different pathways, may operate through common effects on cortical circuits. Overall, our study represents one of few reports of changes in neural responses (both single neurons and populations) accompanied with behavioral improvement after gaining visual experience with an image. This work sheds light on the ways in which accumulated sensory experience can reshape neural activity to impact behavior.

## Acknowledgements

We are grateful to Samantha Schmitt for assistance with data collection, Karen McCracken for animal care, Ben Cowley for assistance with natural image stimulus selection, and Qichao Wu for assistance with spike sorting. PLS was supported by NIH EY031975. MAS was supported by NIH EY029250 and MH118929.

